# Beta-cell mass expansion during puberty involves serotonin signaling and determines glucose homeostasis in adulthood

**DOI:** 10.1101/2022.04.06.487366

**Authors:** Anne-Laure Castell, Mélanie Ethier, Grace Fergusson, Caroline Tremblay, Clara Goubault, Marie Baltz, Dorothée Dal Soglio, Julien Ghislain, Vincent Poitout

## Abstract

Puberty is associated with transient insulin resistance that normally recedes at the end of puberty; however, in overweight children insulin resistance persists leading to an increased risk of type 2 diabetes. The mechanisms whereby pancreatic β cells adapt to pubertal insulin resistance, and how they are affected by the metabolic status, have not been investigated. Here we show that puberty is associated with a transient increase in β-cell proliferation in rats and humans of both sexes. In rats, β-cell proliferation correlated with a rise in growth hormone (GH) levels. Serum from pubertal rats and humans promoted β-cell proliferation, suggesting the implication of a circulating factor. In pubertal rat islets, expression of genes of the GH/serotonin (5-HT) pathway underwent changes consistent with proliferative effect. Inhibition of the pro-proliferative 5-HT receptor isoform HTR2b blocked the increase in β-cell proliferation in pubertal islets ex vivo and in vivo. Peri-pubertal metabolic stress blunted β-cell proliferation during puberty and led to altered glucose homeostasis later in life. This study identifies a role of GH/GHR/5-HT/HTR2b signaling in the control of β-cell mass expansion during puberty and a mechanistic link between pubertal obesity and the risk of developing type 2 diabetes.

## INTRODUCTION

Puberty is a period of considerable dynamic hormonal changes characterized by activation of the hypothalamic–pituitary–gonadal (HPG) axis with subsequent secretion of sex steroids and an increase in growth hormone (GH) release at its highest rate in life. Linked to the action of these hormones, puberty is marked by an accumulation of fat mass and a decrease in insulin sensitivity associated with hyperinsulinemia (1–5). While insulin sensitivity is restored at the end of puberty in normal-weight youth, insulin resistance persists in obese adolescents (6–8). Childhood obesity is a major risk factor for metabolic and cardiovascular complications including type 2 diabetes (T2D) (9–11). In a longitudinal study, Reinehr et al. (12) showed that in the context of obesity, entering into puberty doubles the risk of developing metabolic complications in both males and females. Hence, puberty, much like intrauterine and early postnatal development, appears to be a critical period during which metabolic stress can lead to the development of metabolic disease later in life (13,14). Thus, it is critical to identify the factors contributing to changes in glucose homeostasis during puberty under physiological and pathological conditions of obesity, and to understand how such factors contribute to an increased risk of T2D in young adulthood (15).

Expansion of functional β-cell mass occurs under physiological (e.g. pregnancy) or pathological (e.g. obesity) conditions of insulin resistance which demand a commensurate rise in insulin output from the pancreatic β cells to maintain glucose homeostasis (16,17). In rodents and humans, β-cell mass expansion results from several mechanisms including replication of existing β cells (18). Although limited evidence suggests that β-cell replication increases in humans during puberty (19,20), the mechanisms controlling β-cell proliferation in the face of pubertal insulin resistance remain largely unexplored.

At the onset of puberty, gonadotropin-releasing hormone (GnRH)-expressing neurons in the hypothalamus secrete GnRH, which activates receptors in the anterior pituitary to produce luteinizing hormone (LH) and follicle-stimulating hormone (FSH). LH and FSH then stimulate gonadal steroid secretion of estrogens from the ovaries and testosterone from the testes. In addition to their direct endocrine actions on target tissues, sex steroids stimulate GH production from the pituitary to activate the GH/Insulin-like growth factor-1 (IGF-1) axis (21,22). By binding to the growth hormone receptor (GHR) expressed in the β cell (23), GH is implicated in β-cell mass expansion perinatally and in pregnancy via autocrine/paracrine serotonin ((5-hydroxytryptamine (5-HT)) signaling (24–26). During the perinatal period, GH stimulates 5-HT production in β cells by increasing levels of the 5-HT synthesizing enzyme tryptophan hydroxylase-1 (TPH1) which in turn stimulates β-cell proliferation via the G protein-coupled receptor HTR2b (25). During pregnancy, 5-HT acts downstream of placental lactogens which signal via the prolactin receptor to stimulate β-cell proliferation (26). In the maternal β cell TPH1 and HTR2b expression and 5-HT production are increased, while expression of the inhibitory isoform of the 5-HT receptor HTR1d is reduced. GHR signaling also plays critical roles in regulated insulin secretion and compensatory β-cell proliferation in obese mice (27) and 5-HT acting via HTR2b and HTR3a promotes insulin secretion in rodent and human islets (28,29). However, whether GH/5-HT signaling contributes to β-cell adaptation to puberty has not been established.

The present study was designed to: 1-Determine whether β cells undergo a wave of replication during puberty in rodents and humans; 2-Investigate the role of the GH/GHR/5-HT/HTR2b signaling pathway in controlling β-cell proliferation during puberty; 3-Assess whether high-fat feeding during puberty in rats compromises the β-cell adaptive response and leads to abnormal glucose homeostasis in adulthood.

## RESULTS

### Puberty in rats is associated with glucose intolerance in both sexes

To establish a preclinical model of β-cell compensation during puberty, we characterized glucose homeostasis in male and female rats from weaning to adulthood (Fig. 1 & Supplementary Fig. 1). Based on the clinical evaluation of the secondary sex characteristics (testicular length in male and vaginal opening in female) and testosterone levels we defined the peri-pubertal period in males between 4.5 to 7.5 wk of age and puberty onset in females beginning at 4 wk of age (Fig. 1A-C). During this period, we observed a significant increase in fasting insulinemia in both sexes (Fig. 1D) with stable blood glucose levels (Supplementary Fig. 1A). i.p. glucose tolerance tests (IPGTT) revealed a decrease in glucose tolerance in male (Fig. 1E) and female (Fig. 1F) rats at puberty that persisted at early adulthood (∼8-9 wk of age) in both sexes (Fig. 1E & F). Higher insulin excursions were observed during the IPGTT in male and female rats at puberty compared to weaning or adulthood (Fig. 1G & H). The higher circulating insulin levels at stable blood glucose strongly suggest that glucose intolerance is mostly due to insulin resistance, in line with human data (1–5).

**Figure 1.**
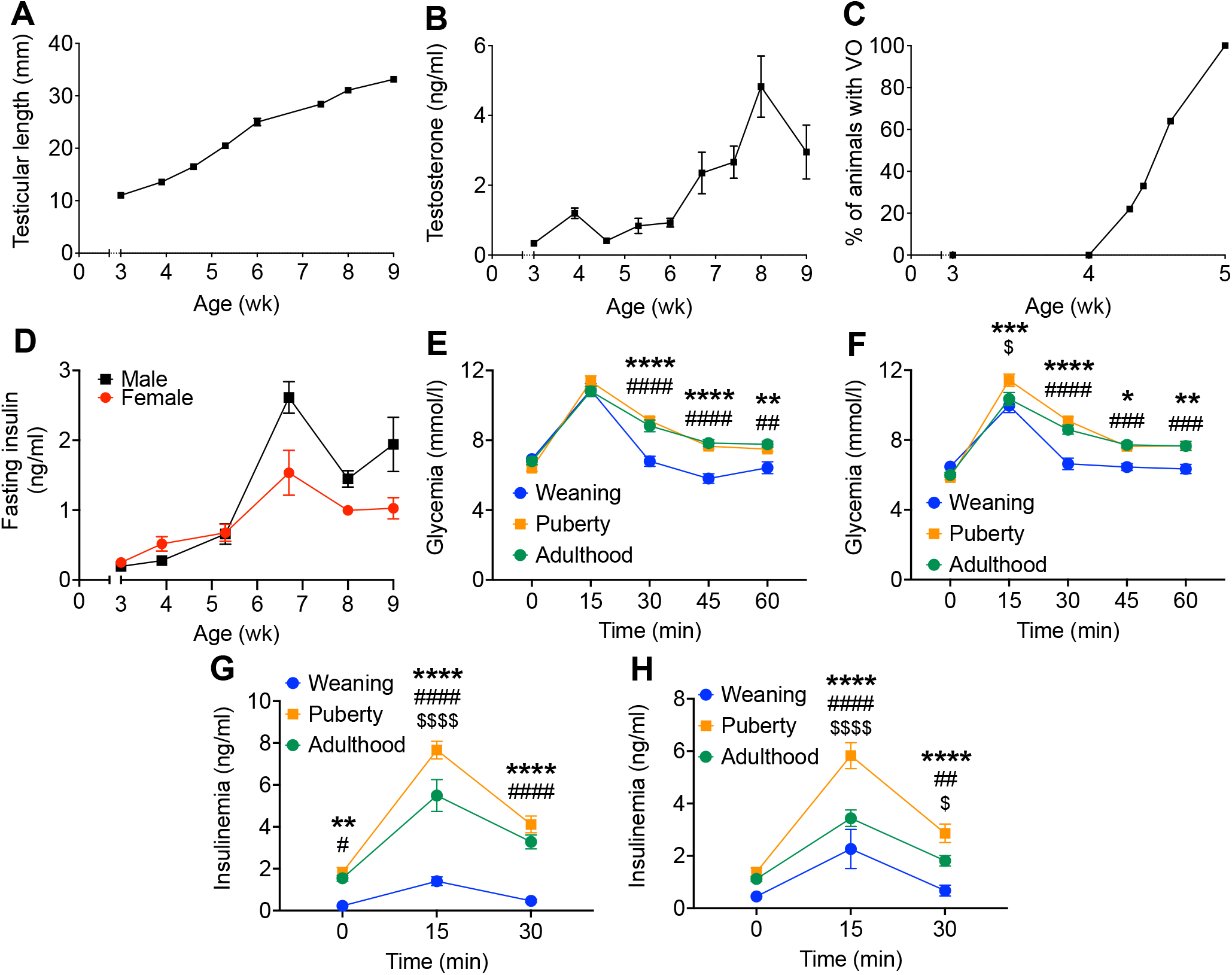
Glucose tolerance is decreased and insulin secretion increased during puberty in female and male rats. (**A**,**B**) Right testicular length (**A**), and plasma testosterone levels (**B**) in male rats. (**C**) Percentage of female rats with vaginal opening (VO). **(D**) Fasting insulin levels in male (black square) and female (red circle) rats (n=6-9). (**E-H**) Glycemia (**E**,**F**) and insulinemia (**G**,**H**) following IPGTT (1g/kg) in male (**E**,**G**) and female (**F**,**H**) rats at weaning (3 wk of age, blue), puberty (∼6 wk of age, orange) or young adulthood (∼9 wk of age, green) (n=10-15). Data are expressed as mean +/-SEM. *p<0.05, **p<0.01, ***p<0.005, ****p<0.001 comparing puberty to weaning group, #p<0.05, ##p<0.01, ###p<0.005, ####p<0.001 comparing the adult to weaning group and $p<0.05, $$$$p<0.001 comparing the puberty to adult group following one-way ANOVA (**E-H**) with Tukey’s multiple comparisons test.

### Beta-cell proliferation increases during puberty in rats and humans

To determine whether the increase in circulating insulin observed during puberty was due to a rise in β-cell mass, we assessed β-cell proliferation and mass by immunostaining of pancreatic sections from male and female rats sequentially from weaning to adulthood (Fig. 2A-E & Supplementary Fig. 1B-C). The onset of puberty in both sexes was marked by a significant and transient increase in the percentage of Ki67^+^/Ins^+^ (Fig. 2A & B) which was confirmed by staining for Nkx6.1 as a complementary β-cell marker (Supplementary Fig. 1B-C). Correspondingly, β-cell mass increased overtime from weaning to adulthood in both sexes (Fig. 2C), which was associated with an overall increase in islet size in males (Fig. 2D), but not females (Fig. 2E). To demonstrate that the increase in β-cell proliferation was effectively triggered by the onset of puberty, male rats were exposed to the GnRH receptor antagonist Cetrorelix (30). Cetrorelix blocked the onset puberty, as shown by the decrease in testicular weight (Fig. 2F) and testosterone levels without changes in IGF1 levels (Supplementary Fig. 1D-E), and also diminished β-cell proliferation from its peak at 5 wk of age (Fig. 2G).

**Figure 2.**
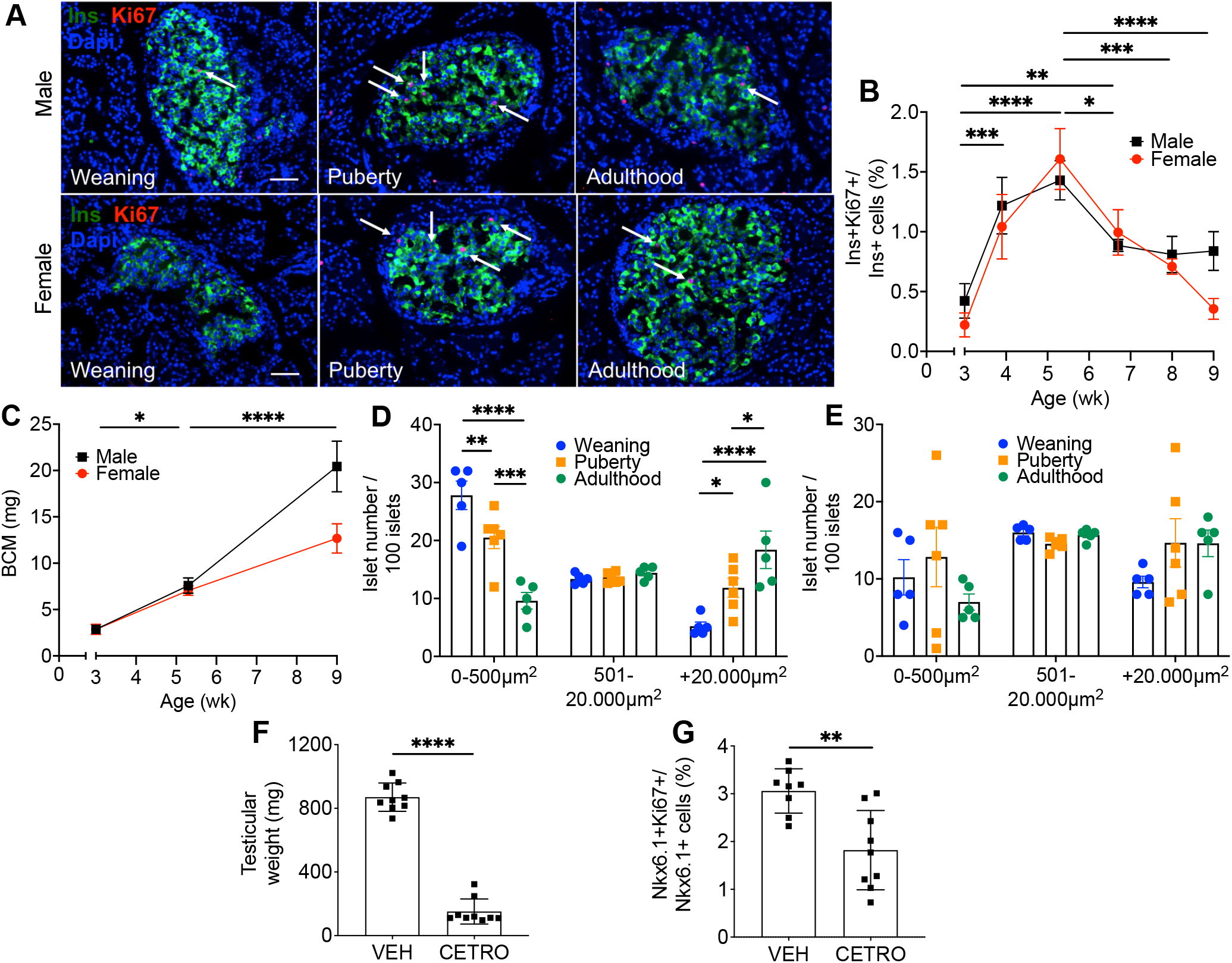
Beta-cell proliferation and mass are increased during puberty in rats and β-cell proliferation is blocked by GnRH antagonism. (**A**,**B**) β-cell proliferation as assessed by immunofluorescent staining of pancreatic sections for Ki67 and insulin (Ins) in male and female rats from 3-9 wk of age. (**A**) Representative sections showing Ins (green), Ki67 (red) and nuclei (Dapi, blue) from male (top) and female (bottom) rats at weaning (3 wk-old), puberty (∼5 wk-old) or young adulthood (9 wk-old). Arrows show positive nuclei for Ki67. Scale bars, 50μm. (**B**) β-cell proliferation as a percentage of Ki67+ insulin+ cells over insulin+ cells in males (black square) (n=3-5) and females (red circle) (n=3-5). (**C**) β-cell mass (BCM) in males (black square) (n=5-6) and females (red circle) (n=5-6). (**D**,**E**) Islet distribution by size in male (**D**) and female (**E)** rats at weaning (blue circle), puberty (orange square) or adulthood (green circle). (**F**,**G**) Male rats treated with Cetrorelix (CETRO; 100μg/d) or vehicle (VEH) from D25 to D37 (n=8-9). Right testicular weight (**F**) and β-cell proliferation (**G**) were assessed at D38. β-cell proliferation was measured by immunofluorescent staining of pancreatic sections for Ki67 and Nkx6.1 and presented as a percentage of Ki67+ Nkx6.1+ cells over Nkx6.1+ cells. Data represent individual or mean values and are expressed as mean +/-SEM. *p<0.05, **p<0.01, ***p<0.005, ****p<0.001 following one-way (**B**,**C**) or two-way (**D**,**E**) ANOVA with Tukey’s multiple comparisons test or compared to the VEH group (**F**,**G**) following unpaired Student’s t-test.

In post-mortem pancreatic samples from 8-15 yr-old children (5 males and 8 females) we detected several Ki67^+^/C-pep^+^ cells in all samples at pubertal Tanner stages III-IV (corresponding to the peak of puberty (31)) both in males (Fig. 3C & D) and females (Fig. 3B & D), but none in pre- and onset-(Tanner stages I-II) or post-puberty (Tanner stage V) donors (Fig. 3A & D).

**Figure 3.**
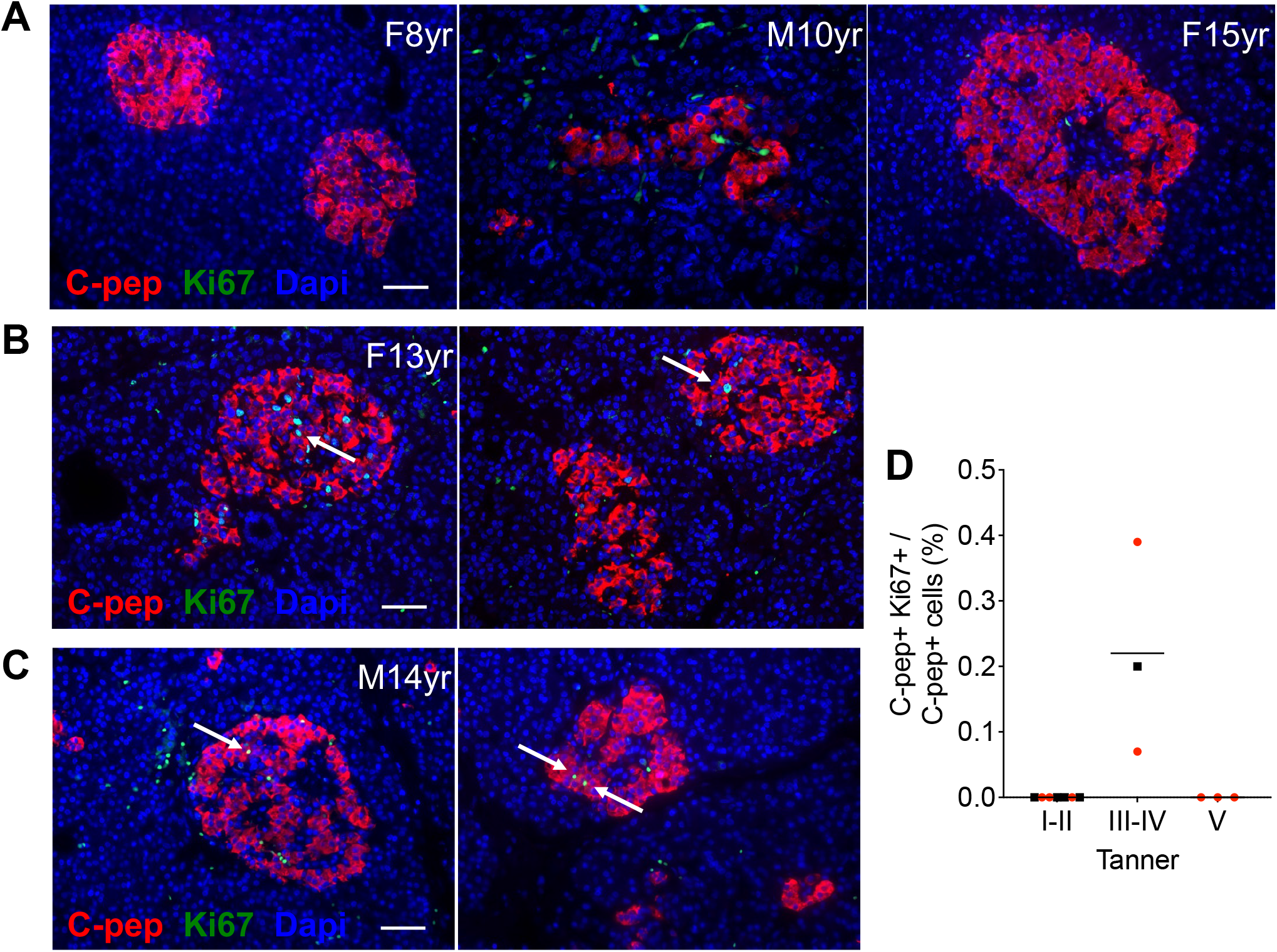
Beta-cell proliferation is increased during puberty in humans. β-cell proliferation in post-mortem pancreatic sections of male (M) and female (F) children from 8 to 15 yr-old as assessed by immunofluorescent staining for Ki67 and C-peptide (C-pep). (**A-C**) Representative sections showing C-pep (red), Ki67 (green) and nuclei (Dapi, blue) from Tanner I-II (8 yr-old female and 10 yr-old male) and V (15 yr-old female) (**A**), and Tanner III-IV (13 yr-old female and 14 yr-old male) (**B**,**C**), as indicated. Arrows show positive nuclei for Ki67. Scale bars, 50μm. (**D**) β-cell proliferation presented as a percentage of Ki67^+^ C-pep^+^ cells over C-pep^+^ cells grouped according to Tanner stage in males (black square) (n=5) and females (red circle) (n=8).

### Pubertal serum stimulates β-cell proliferation in rat and human islets

To investigate whether β-cell proliferation at puberty onset is mediated by a circulating factor, we exposed pubertal rat islets to decomplemented weaning or pubertal rat serum for 72 h and measured islet cell proliferation by flow cytometry (Fig. 4A). Pubertal serum significantly increased α- and β-cell proliferation compared to serum collected at weaning (Fig. 4A-C). Accordingly, decomplemented human serum from children > 11 yr old, but not children < 8 yr old, increased β-cell proliferation in sex-matched adult human islets to a similar level as the DYRK1A inhibitor harmine used as a positive control (32) (Fig. 4D). Interestingly, unlike in rat islets (Fig. 4C) human α cells did not respond to pubertal serum (Fig. 4E).

**Figure 4.**
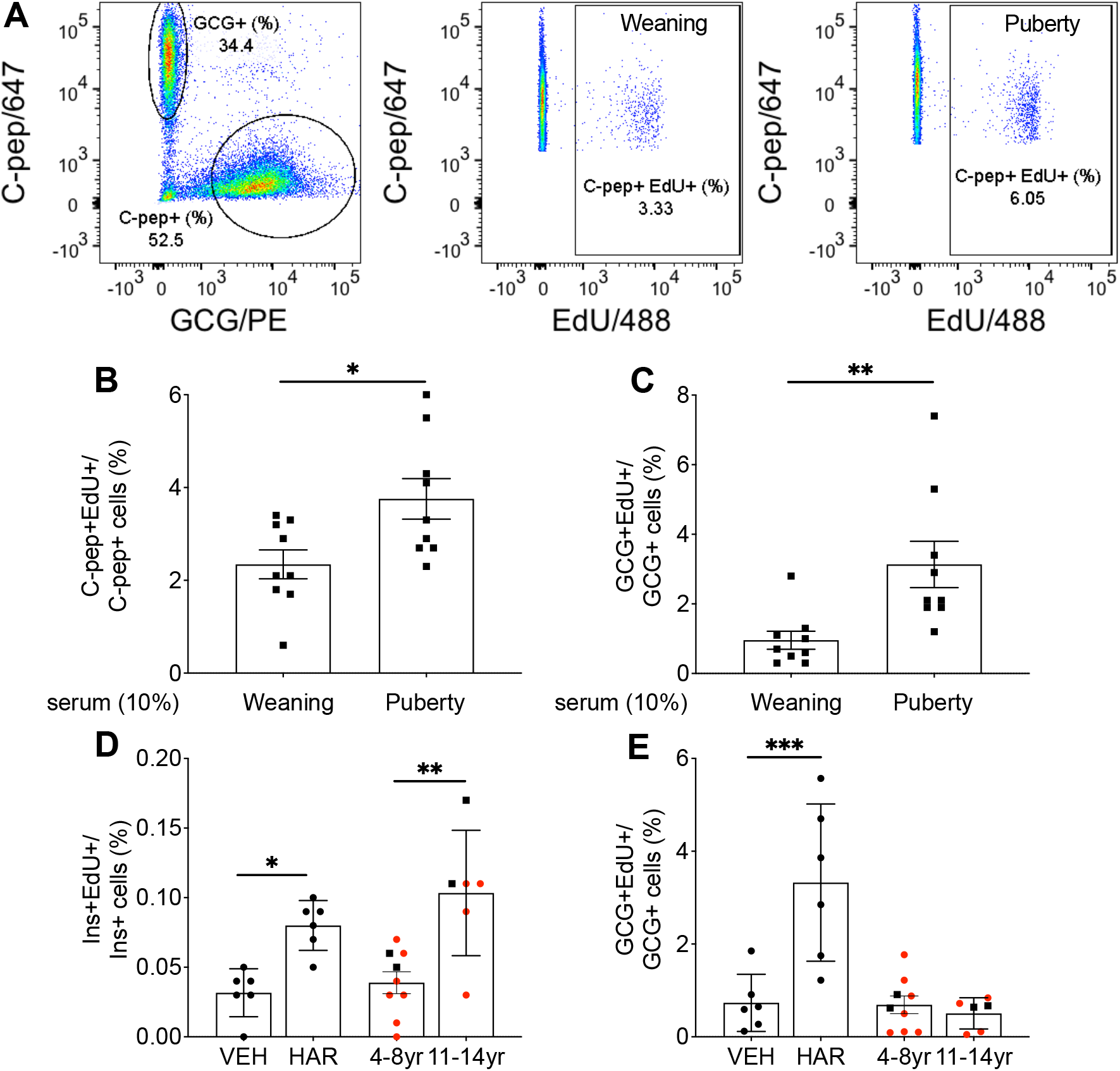
Pubertal but not pre-pubertal serum stimulates β-cell proliferation in isolated rat and human islets. (**A-C**) Male rat islets isolated at puberty were exposed to pre-pubertal (Weaning) or pubertal (Puberty) rat serum for 72h and β- and α-cell proliferation assessed by flow cytometry following staining for EdU and C-peptide (C-pep) or glucagon (GCG), respectively. (**A**) Representative plots showing gates used to select C-pep^+^ and GCG^+^ cells and EdU^+^/C-pep^+^ cells following exposure to weaning or pubertal serum as indicated. β-(**B**) and α-(**C**) cell proliferation as a percentage of EdU^+^ C-pep^+^ or EdU^+^ GCG^+^ cells over C-pep^+^ or GCG^+^ cells. (**D**,**E**) Adult human islets were exposed to Harmine (HAR, 10μM) or to sex-matched pre-pubertal (4-8yr) or pubertal (11-14yr) human male (black square) or female (red circle) serum for 72h and β- and α-cell proliferation assessed by flow cytometry following staining for EdU and insulin (Ins) or GCG, respectively. β-(**D**) and α-(**E**) cell proliferation presented as a percentage of EdU^+^Ins^+^ or EdU^+^GCG^+^ cells over Ins^+^ or GCG^+^ cells. Data represent individual values and are expressed as mean +/-SEM. *p<0.05 **p<0.01 ***p<0.005 following unpaired Student’s t-test (**B**,**C**) or following one-way ANOVA with Tukey’s multiple comparisons test (**D**,**E**). VEH, vehicle.

Taken together, these data suggest that β-cell proliferation during puberty in rodents and humans is triggered by a circulating factor.

### HTR2b signaling is implicated in β-cell proliferation during puberty

GH levels increase during puberty in humans (22) and GHR signaling stimulates β-cell proliferation via autocrine/paracrine 5-HT/HTR2b signaling (24–26). Hence, we asked whether this pathway mediates β-cell proliferation during puberty in rats. First, in line with studies in humans, we detected an increase in circulating IGF-1 levels (a surrogate for GH) in both sexes during puberty, concomitant with the rise in β-cell proliferation (Fig. 5A). Second, a membrane-filtered pubertal serum fraction ≥ 3 kDa containing GH stimulated β-cell proliferation, whereas the fraction ≤ 3 kDa depleted in GH was inactive (Supplementary Fig. 2A & B). Third, mRNA expression of GHR, TPH1 and the stimulatory 5-HT receptor isoform HTR2b was increased in islets at the onset of puberty, whereas that of the inhibitory 5-HT receptor isoform HTR1d was diminished (Fig. 5B). Fourth, the selective HTR2b antagonist SB204741 (33) blocked pubertal serum-induced β-cell proliferation in isolated rat islets (Fig. 5C). Finally, to investigate the contribution of HTR2b signaling in vivo we administered SB204741 to male rats from D25 to D37 and assessed glucose tolerance and β-cell proliferation at D38 (Fig. 5D). Body weight was similar in both SB204741- and vehicle-treated groups (Supplementary Fig. 3A). During the IPGTT, animals treated with SB204741 showed higher glycemia at 15 min (Fig. 5E) and higher area under the glucose curve during the first 30 minutes of the test (Supplementary Fig. 3B) suggesting mild glucose intolerance. Insulinemia was not significantly different between the groups (Fig. 5F; Supplementary Fig. 3C), yet the number of BrdU^+^ β cells was lower in the SB204741-injected group at D38 (Fig. 5G).

**Figure 5.**
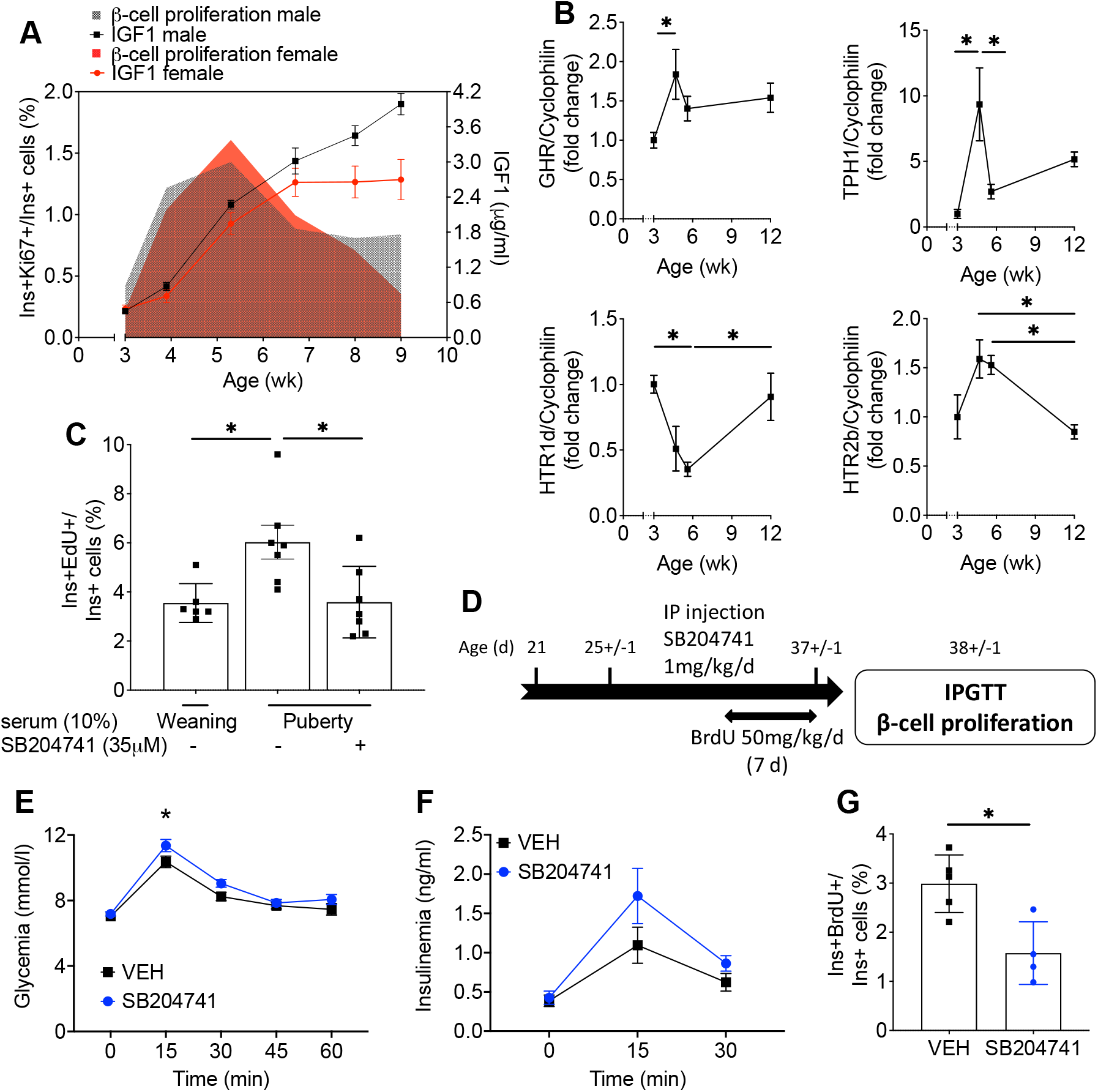
Beta-cell proliferation during puberty correlates with GH-serotonin pathway gene expression and is blocked by HTR2b antagonism in rat islets ex vivo and in vivo. (**A**) Plasma IGF1 levels and β-cell proliferation (from Figure 2B) in female (red) and male (black) rats from 3-9 wk of age (n=4-7). (**B**) GHR, TPH1, HTR1d and HTR2b mRNA levels in rat islets isolated at different ages as indicated (n=4-5). mRNA was quantified by RT-PCR and normalized to cyclophilin. Data are presented as the fold change over the pre-pubertal time point (3 wk of age). (**C**) Male rat islets isolated at puberty were exposed to 10% weaning (3 wk-old) or pubertal (∼5 wk-old) rat serum in the presence of SB204741 (35μM) for 72h and β-cell proliferation assessed by flow cytometry following staining for EdU and Ins and presented as a percentage of EdU^+^Ins^+^ over Ins^+^ cells (n=6-7). (**D**) Male rats were exposed to SB204741 (1mg/kg/d) or vehicle (VEH) from D25 to D37 and BrdU from D30-37 after which IPGTT (1g/kg) were performed and β-cell proliferation assessed by immunofluorescent staining of pancreatic sections for BrdU and Ins. Glycemia (**E**) and insulinemia (**F**) during the IPGTT in the SB204741-injected group (blue) and the control group (VEH, black). (**G**) β-cell proliferation as a percentage of BrdU^+^Ins^+^ over Ins^+^ cells (n=4-5). Data are expressed as mean +/-SEM. *p<0.05 following one-(**B**,**C**) or two-(**E**,**F**) way ANOVA with Tukey’s multiple comparisons test or following unpaired Student’s t-test (**G**).

Taken together, these data suggest that HTR2b signaling is implicated in β-cell proliferation during puberty.

### Peri-pubertal high-fat diet is associated with glucose intolerance and reduced β-cell mass in adulthood

To assess the impact of peri-pubertal metabolic stress on glucose homeostasis, male rats were exposed to high-fat diet (HFD) from 4 to 8 wk of age followed by normal chow diet until 12 wk of age (HFD-CHOW), or chow diet throughout (CHOW-CHOW) (Fig. 6A). Body weight and cumulative food intake increased in the HFD-CHOW group compared to controls beginning at 8 wk of age (Supplementary Fig. 4A & B). IPGTT performed at 8 wk of age revealed glucose intolerance with higher insulinemia in the HFD-fed group compared to controls (Fig. 6B & C). Interestingly, despite the switch to chow diet at 8 wk the animals that had been fed HFD from wk 4 to 8 were still glucose-intolerant at 12 wk of age (Fig. 6D) without an increase in insulin levels during the test (Fig. 6E), suggesting defective insulin secretion. To further explore a potential β-cell secretory defect we performed hyperglycemic clamps (HGC) on HFD-CHOW and CHOW-CHOW groups at 12 wk of age. During the steady state of the clamp (50 to 80 min) blood glucose levels were in the target range in both groups (Supplementary Fig. 4C). The glucose infusion rate (GIR) was lower in the HFD-CHOW group (Fig. 6F), indicative of insulin resistance, and C-peptide levels trended lower (Fig. 6G), consistent with defective insulin secretion.

**Figure 6.**
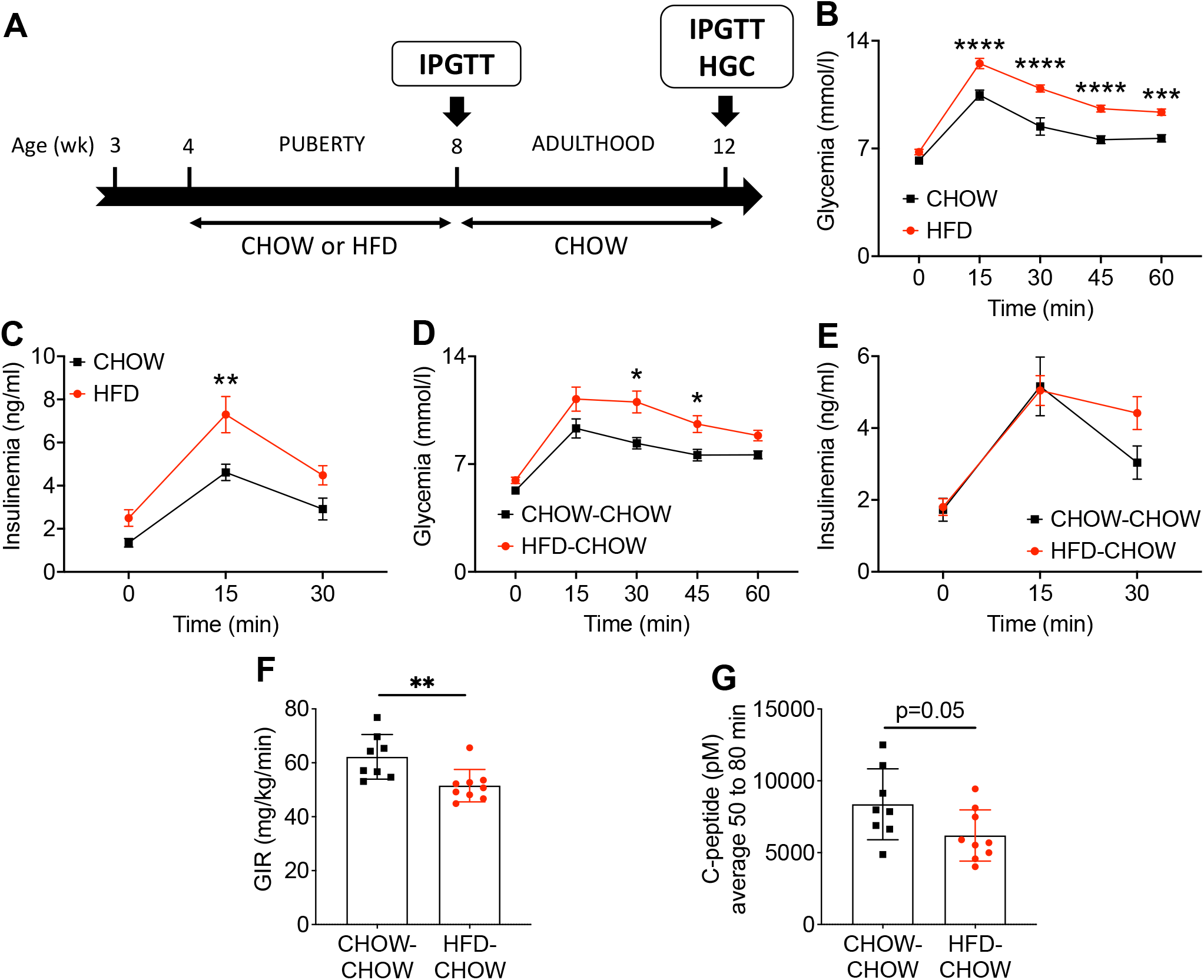
HFD during puberty leads to glucose intolerance in adulthood. IPGTT and HGC were performed on male rats fed a HFD (HFD, red) or a chow diet (CHOW, black) from 4 to 8 wk of age followed by a switch to a chow diet for both groups (HFD-CHOW, red; CHOW-CHOW, black) until 12 wk of age (**A**). Glycemia (**B**,**D**) and insulinemia (**C**,**E**) during the IPGTT (1g/kg) performed at 8 (n=7-9) (**B**,**C**) and 12 (n=6-9) (**D**,**E**) wk of age. Glucose infusion rate (GIR) (**F**) and C-peptide levels (**G**) during the HGC performed at 12 wk of age (n=8-9). Data are expressed as mean +/-SEM. *p<0.05, **p<0.01, ***p<0.005, ****p<0.001 following two-way ANOVA with Sidak’s multiple comparisons test (**B-E**) or unpaired Student’s t-test (**F**,**G**).

To assess whether HFD during puberty impacts β-cell mass expansion, we measured β-cell proliferation and mass in HFD-CHOW and CHOW-CHOW groups (Fig. 7A). β-cell proliferation was decreased at 5 (Fig. 7B & C) and 8 (Fig. 7D & E) wk of age in HFD-fed animals compared to the CHOW-fed group. Consequently, β-cell mass was reduced at 12 wk of age in the HFD-CHOW group (Fig. 7F).

**Figure 7.**
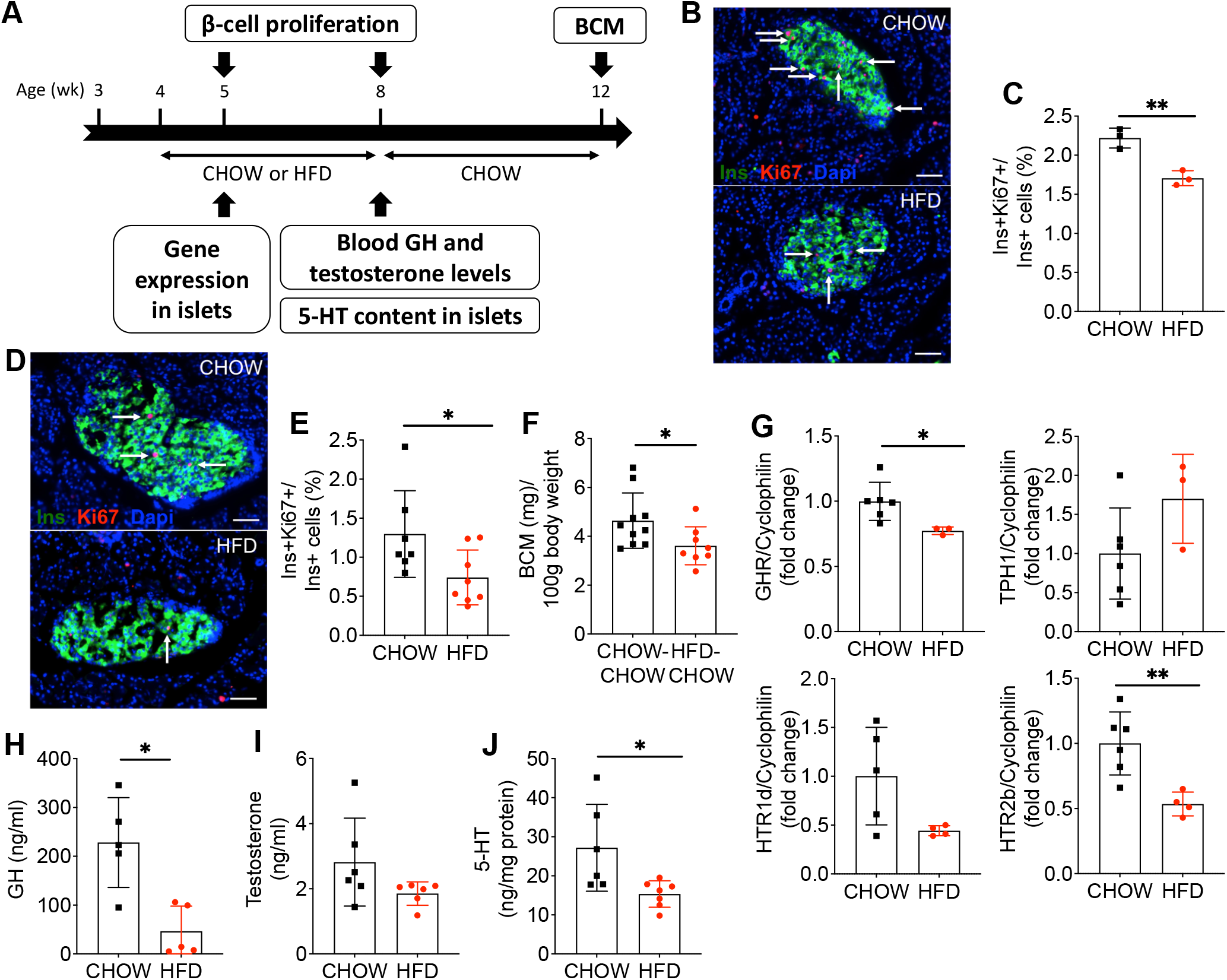
HFD during puberty mitigates β-cell proliferation and GH-serotonin pathway hormone levels and gene expression and reduces β-cell mass in adulthood. β-cell proliferation and mass, plasma hormone and islet gene expression and 5-HT content were assessed in male rats fed a HFD (HFD, red) or a chow diet (CHOW, black) from 4 to 8 wk of age followed by a switch to a chow diet for both groups (HFD-CHOW, red; CHOW-CHOW, black) until 12 wk of age (**A**). β-cell proliferation was assessed at 5 (**B**,**C**) and 8 (**D**,**E**) wk of age by immunofluorescent staining of pancreatic sections for Ki67 and Ins. (**B**,**D**) Representative sections showing Ins (green), Ki67 (red) and nuclei (Dapi, blue). Arrows show positive nuclei for Ki67. Scale bars, 50μm. (**C**,**E**) β-cell proliferation as a percentage of Ki67^+^Ins^+^ cells over Ins^+^ cells (n=3-8). (**F**) β-cell mass (BCM) as a fraction of body weight at 12 wk of age (n=8-10). (**G**) GHR, TPH1, HTR1d and HTR2b mRNA levels in islets isolated from rats at 5 wk of age (n=3-6). mRNA was quantified by RT-PCR and normalized to cyclophilin. Data are presented as the fold change over the CHOW diet group. Plasma GH (**H**) and testosterone (**I**) levels and islet 5-HT content normalized to total protein (**J**) in rats at 8 wk of age (n=6-7). Data represented individual values and are expressed as mean +/-SEM. *p<0.05, **p<0.01 following unpaired Student’s t-test as compared to the control CHOW or CHOW-CHOW group.

Overall, these data indicate that pubertal HFD blunts the normal β-cell proliferative response and leads to glucose intolerance and lower β-cell mass in adulthood.

### Peri-pubertal HFD dampens GH/5-HT/HTR2b signaling in islets

Given the role of HTR2b in pubertal β-cell proliferation shown in Fig. 5, we tested whether this pathway was affected by HFD (Fig. 7). One wk after initiation of HFD, GHR and HTR2b mRNA expression was decreased compared to controls while TPH1 and HTR1d were similar between groups (Fig. 7G). Circulating GH levels were markedly lower after 4 wk of HFD (Fig. 7H) while testosterone levels were unchanged (Fig. 7I). Accordingly, 5-HT concentrations in islets were lower in the HFD group (Fig. 7J).

## DISCUSSION

The objectives of this study were to identify the mechanisms of β-cell adaptation to insulin resistance during puberty and to ascertain whether this process is perturbed by metabolic stress. We showed that puberty is associated with a transient increase in β-cell proliferation mediated by a circulating factor in rats and humans of both sexes. Mechanistically, pubertal islets undergo a phenotypic switch in the expression of genes of the GH/5-HT signaling pathway that favors β-cell proliferation, and inhibition of HTR2b signaling blocks the β cell response ex vivo and in vivo. Finally, peri-pubertal HFD blunts 5-HT signaling and the physiological rise in β-cell proliferation, leading to impaired β-cell mass and glucose homeostasis later in life.

Our observations are consistent with previous study in neonatal male Sprague Dawley rats describing an increase in β-cell mass at D24 and D31 associated with an increase in β-cell proliferation (34). In human samples, we detected proliferating β cells at mid-puberty (Tanner stages III-IV). Accordingly, Lam et al (20) observed an increase in islet endocrine cell proliferation in adolescents and young adults, although the proliferating islet cells did not express β-cell markers contrary to our observations. Interestingly, the increase in β-cell proliferation we detected in human samples mirrored the peak of physiological insulin resistance that also occurs at Tanner stages III-IV and returns to pre-pubertal levels at Tanner stage V (1,3,4,35). Overall, the data indicate that β cells undergo a wave of proliferation in response to pubertal insulin resistance in rodents and humans.

Our observation that pubertal serum from rats and humans stimulates β-cell proliferation demonstrates the involvement of a circulating factor. Importantly, decomplemented serum was completely devoid of insulin (data not shown), excluding the possibility that β-cell replication is directly stimulated by circulating insulin. In addition, the increase in β-cell proliferation preceded the peak in plasma insulin levels in both sexes in rats. The reduction in β-cell proliferation following administration of the GnRH receptor antagonist Cetrorelix in rats supports a role of the HPG axis in β-cell mass expansion during puberty. Gonadotrophins released downstream of GnRH stimulate the secretion of testosterone and estrogens which, in turn, trigger GH secretion from the pituitary. A possible role for sex hormones in β-cell adaptation to puberty is suggested by the observation that castration of young adult male rats leads to a reduction in β-cell mass which is reversed by testosterone administration (36). Accordingly, testosterone and its metabolites acting via the androgen and estrogen receptors are established regulators of β-cell function and mass in males (37,38). However, our attempts to demonstrate a direct role of testosterone in pubertal serum-induced β-cell proliferation were unsuccessful (data not shown). GH is another potential candidate circulating factor that could trigger β-cell proliferation in puberty. Indeed, circulating levels of IGF-1, a surrogate for GH, increased concomitantly with β-cell proliferation during puberty in rats. In serum fractionation studies, only the GH-containing fraction (greater 3 kDa) retained the ability to stimulate β-cell proliferation. Similarly in humans, GH/IGF-1 secretion are at their highest levels during puberty (39). Furthermore, GH causes insulin resistance (40–42) and pubertal insulin resistance positively correlates with GH and IGF-1 levels (43,44). GH acting through GHR increases β-cell proliferation in islets ex vivo (45–48) and conversely, whole body and β-cell specific GHR deficiency in mice are associated with a decrease in β-cell proliferation and mass (25,27,49,50). Overall, our data suggest a role for GH in promoting β-cell proliferation in puberty that remain to be formally confirmed by identification of the circulating factor mediating this effect.

At the islet level, we observed changes in mRNA expression similar to the pattern of expression of maternal islets during pregnancy (26) and suggestive of a phenotype consistent with 5-HT-induced proliferation, namely increased TPH1 and HTR2b and decreased HTR1d. Accordingly, the HTR2b antagonist SB204741 blocked the increase in β-cell proliferation in isolated rat islets in response to pubertal rat serum and abrogated pubertal β-cell proliferation in vivo in rats. Hence, similar to the perinatal period (25) and pregnancy (26), 5-HT signaling controls β-cell proliferation during puberty. This model predicts that 5-HT is synthesized in β cells in response to GHR activation and acts in an autocrine/paracrine fashion, and is therefore entirely consistent with the possibility that GH is the circulating factor mediating this response.

Peri-pubertal HFD in rats blunted the β-cell proliferative response and reduced β-cell mass in adulthood despite a 4-wk “wash-out” period during which the animals received a chow diet. The decrease in β-cell mass was associated with impaired glucose tolerance and insulin secretion. In agreement with our findings, Holtrup et al. (51) found that mice fed a HFD during the peri-pubertal period displayed glucose intolerance and insulin resistance in adulthood. Glucose homeostasis was also more severely impaired in rats fed a HFD before adulthood compared to those receiving HFD only during adulthood (52). In contrast, in a recent study Glavas et al. (53) did not detect impaired glucose homeostasis or changes in β-cell mass when mice were fed a HFD exclusively during the peri-pubertal period, which may have been due to lower fat content of the diet compared to the present study. Nevertheless, increased diabetes incidence was observed when HFD was initiated in mice at pre-weaning and peri-pubertal compared to post-pubertal stages (53), consistent with a more severe outcome following high-fat feeding early in life. Interestingly, resistance to metabolic impairment in mice fed a HFD post-weaning is associated with higher circulating IGF-I levels and lean mass (54). In contrast, although IGF-1 levels were unchanged by HFD in our studies (data not shown), we observed that pubertal HFD reduced plasma GH levels, consistent with an effect of metabolic stress on GH levels in human (55,56). Pubertal HFD also reduced GHR gene expression in islets, and decreased islet 5-HT content and HTR2b gene expression. 5-HT contributes to β-cell compensation to metabolic stress as defects in insulin secretion and glucose tolerance are observed in β-cell specific TPH1 and 5-HT receptor HTR3a KO mice under HFD (28). Taken together with our data supporting a role of 5-HT in β-cell proliferation at puberty, we propose that pubertal HFD leads to reduced 5-HT signaling with resulting impairment of functional β-cell mass in adulthood. Hence, much like the perinatal period (25), pubertal β-cell replication likely contributes to adult β-cell mass and perturbations during this period have lasting detrimental effects on glucose homeostasis.

Because of the rising prevalence of childhood obesity, prediabetes and T2D have become increasingly common in youth (57,58). Puberty is a critical window for the establishment of metabolic health such that physiological changes during this period influence diabetes risk (12,14). In particular, pubertal insulin resistance is exacerbated in obese adolescents and does not return to pre-pubertal values (6,7). However, adolescent obesity is also associated with reduced β-cell function which may augment diabetes risk (8,59,60). Hence, much like during development or early postnatal life the pubertal pancreatic islet may be a key target for metabolic programming that affects risk of metabolic disease later in life (13,60).

In conclusion, this study provides evidence for a role of GH/GHR/5-HT/HTR2b signaling during puberty in the control of β-cell proliferation and mass and sheds light on the molecular pathways linking puberty and obesity to the risk of developing T2D later in life.

## RESEARCH DESIGN AND METHODS

### Reagents

The GnRH receptor antagonist Cetrorelix was from Cayman Chemicals (#23910-10; Ann Arbor, MI, USA), the selective HTR2b receptor antagonist SB204741 was from Tocris (#1372/10; Toronto, ON, Canada); BrdU was from Sigma Aldrich (#B5002; Saint-Louis, MI, USA); The Amicon Ultra-0.5 Centrifugal Filter Units were from Millipore (#UFC500324; Burlington, MA, USA).

### Animals

A Wistar rat (Charles River Laboratories, Saint-Constant, QC, Canada) colony was maintained in-house. Animals were housed under controlled temperature on a 12 h light–dark cycle with free access to water and fed ad libitum with normal chow diet (CHOW; #2018 Teklad Global 18% protein rodent diet; 58% carbohydrate, 24% protein and 18% fat on a caloric basis; Harlan Teklad, Madison, WI) or HFD (#D12492; 60% fat, 20% protein and 20% carbohydrate on a caloric basis; Research Diets Inc., New Brunswick, NJ). The animals received HFD or CHOW diet from 4 to 8 wk of age, and then CHOW diet until 12 wk of age. Body weight, food intake and fed blood glucose were monitored weekly during the HFD study. Clinical parameters (right testicular length in males using Vernier caliper; weight and vaginal opening in females), metabolic tests, β-cell proliferation and mass were assessed at different time points from weaning to adulthood as described in the Figure legends. Plasma insulin, C-peptide, testosterone, GH and IGF-1 were measured by ELISA (Alpco Diagnostics; #80-INSRT-E01, #80-CPTRT-E01, #55-TESMS-E01, #22-GHOMS-E01, #22-IG1MS-E01 respectively, Salem, NH, USA).

### In vivo injections

Male rats were injected i.p. from postnatal day (D) 25+/-1 to 37+/-1 with Cetrorelix (100µg/day in 0.4ml of 0.9% of NaCl), SB204741 (1 mg/kg/day in PEG400) or vehicle. At the end of the treatment, body and testicular weight, plasma testosterone and IGF1 levels and β-cell proliferation in pancreatic sections were measured. BrdU (50 mg/kg/day) was injected IP during the last 7 days of treatment. IPGTT and β-cell proliferation were assessed at D38+/-1.

### Metabolic tests

IPGTT were performed on 4-5 hour fasted rats by measuring tail blood glucose (glucometer Accu-Chek, Roche, Indianapolis, IN) at t=0, 15, 30, 45, 60 min and plasma insulin levels at t=0, 15, 30 min after i.p. dextrose administration (1g/kg).

One-step HGC were performed in conscious, ad libitum fed animals as described (62). Briefly, a 20% dextrose solution (Baxter, Mississauga, ON) was infused via a jugular catheter. Rats initially received a 90-second bolus (140 mg/kg/min) and then the glucose infusion rate (GIR) was adjusted to maintain blood glucose between 15.5 and 17.7 mmol/l for 80 min. Blood samples were collected from an arterial catheter to measure glucose, plasma insulin and C-peptide at the time points indicated in the Figure legends.

### Measurement of β-cell proliferation and mass in pancreatic sections

Pancreata were fixed for 4 h in 4% paraformaldehyde (PFA) and cryoprotected overnight in 30% sucrose. Tissue was then embedded in optimal cutting temperature (OCT) compound, frozen, sectioned at 8 μm, and mounted on Superfrost Plus slides (Life Technologies Inc., Burlington, ON, Canada). Antigen retrieval was performed using sodium citrate buffer (pH=6). Beta-cell proliferation was measured by immunofluorescent staining for Ki67, or BrdU and insulin (Ins) or Nkx6.1 as described (63). Primary antibodies and dilutions are listed in Supplementary Table 1. Secondary antibodies were from Jackson ImmunoResearch (West Grove, PA, USA). Images were acquired with a fluorescence microscope (Zeiss, Thornwood, NY). Beta-cell proliferation was expressed as a percentage of double-positive Ki67^+^ (or BrdU^+^) and Ins^+^ (or Nkx6.1^+^) cells over the total Ins^+^ (or Nkx6.1^+^) cells. At least 1,500 β cells were manually counted per condition. The experimenter was blind to group assignments. Beta-cell mass was measured on paraffin sections prepared as described (64) using an anti-insulin antibody (Supplementary Table 1).

### Immunofluorescent staining of human pancreatic sections

Formalin fixed, paraffin embedded sections were deparaffinized and rehydrated. Antigen retrieval was performed using sodium citrate buffer (pH=6). Beta-cell proliferation was measured by immunofluorescent staining for Ki67 and C-peptide and expressed as a percentage of double-positive Ki67^+^ and C-pep^+^ cells over the total C-pep^+^ cells as described above for rats. Primary antibodies and dilutions are listed in Supplementary Table 1. Samples were grouped according to Tanner stage. When the Tanner stage was unknown, age was used to define the pubertal stage (31).

### Rat and human serum

Blood was collected from male rats following abdominal aorta puncture at weaning (D21) or during puberty (D37-D39) and centrifuged at 2,500 rpm for 10 min at 4°C in BD Vacutainer SST tubes (VWR, Mississauga, ON, Canada, #367985). Pre-pubertal (4-8 yr old) or pubertal (11-14 yr old) sera from boys and girls were obtained from Innovative Research Inc (MI, USA). All sera were decomplemented by heating at 56+/-2°C during 30 min prior to use.

### Rat islet isolation

Peri-pubertal (D37+/-1) rat islets were isolated from male rat by collagenase digestion and dextran density gradient centrifugation as described previously (65) and allowed to recover overnight in RPMI-1640 supplemented with 10% (v/v) FBS (Life Technologies Inc, Burlington, ON, Canada), 100 U/ml penicillin/streptomycin (Multicell Wisent Inc, Saint-Jean-Baptiste, QC, Canada) and 11.1 mM glucose prior to use.

### Human islets

Upon reception human islets were maintained in cGMP Prodo Islet Media (Standard; Prodo Laboratories, Aliso Viejo, CA, USA) with 5% (vol./vol.) human AB serum and 1% (vol./vol.) glutamine/glutathione mixture prior to use.

### Rat and human islets culture

Batches of 200 peri-pubertal male rat islets (D37+/-1) were cultured in RPMI-1640 in the presence of 7.5 mM glucose for 4 h prior to the addition of rat serum (10% v/v) for an additional 72 h. Adult human islets were cultured in Prodo media (5.8 mM glucose) with decomplemented sex-matched human serum (10% v/v) for 72 h. SB204741 (35 µM) and 5-ethynyl-2’-deoxyuridine (EdU, 10 µM) was added throughout. Media were replaced daily.

### Beta-cell proliferation in isolated islets

Following treatment islets were dissociated into single cells with accutase (1 μl/islet; Innovative Cell Technologies Inc., San Diego, CA) for 10 min at 37° and β-cell proliferation assessed by flow cytometry as described previously (66). Dead cells were labeled using the LIVE/DEAD™ Aqua (405 nm) (BD Bioscience). EdU was detected using Click-iT Plus EdU Alexa Fluor 488. Immunostaining was performed according to the manufacturer’s instructions (Thermo Fisher Scientific, Waltham, MA). Fluorophore-coupled primary antibodies and dilutions are listed in Supplementary Table 1. Flow cytometry was carried out using an LSRIIB flow cytometer with FACSDiva software (BD Biosciences). Data was analyzed using FACSDiva or FlowJo v10.7 software (Ashland, OR; https://www.flowjo.com/solutions/flowjo). Dead-cell stain, EdU, Ins and glucagon (GCG) labelled cells were detected using the 405-, 488-, 640- and 561-nm lasers coupled with 525/50-, 530/30-, 670/14- and 586/15-nm BP filters, respectively. Proliferation was calculated as the percentage of double-positive EdU^+^ and Ins^+^ cells over the total Ins^+^ cell population. At least 2,300 Ins^+^ cells were counted in each sample.

### 5-HT content in islets

Following isolation, rat islets were allowed to recover in RPMI-1640 supplemented with 10% (v/v) FBS at 11.1 mM glucose overnight, after which islets were lysed with radioimmunoprecipitation assay (RIPA) buffer (0.5 μl/islet), sonicated, and centrifuged at 12,000 x g for 5 min at 4°C. 5-HT was measured in supernatants using an ELISA kit (LDN, Nordhorn, Germany). Total protein content was measured using the BCA assay (Thermo Fisher Scientific). 5-HT content was normalized to total protein.

### Quantitative RT-PCR

Total RNA was extracted from 150 to 200 islets after an overnight recovery in RPMI-1640 supplemented with 10% (v/v) FBS at 11.1 mM glucose using the RNeasy Micro kit (Qiagen, Valencia, CA). RNA was quantified by spectrophotometry using a NanoDrop 2000 (Life Technologies Inc.) and reverse transcribed (1 μg). Real-time PCR was performed using the Rotor-Gene SYBR Green PCR kit (Qiagen). Results are expressed as the ratio of target RNA to cyclophilin A RNA levels and normalized to control islets. Primer sequences are listed in Supplementary Table 2.

### Statistical analyses

Data are expressed as means ± SEM. Statistical analyses were performed using Student’s t test, one-or two-way ANOVA with Tukey’s, Sidak’s or Dunnett’s post hoc test adjustment for multiple comparisons, as appropriate, using GraphPad Prism 9 Software (San Diego, CA). p<0.05 was considered significant.

### Study approval

All the animal studies were approved by the Institutional Committee for the Protection of Animals at the Centre de Recherche du Centre Hospitalier de l’Université de Montréal (CRCHUM). Post-mortem pancreatic paraffin sections from children 8-15 yr old (5 males and 8 females) were provided by the Alberta Diabetes Institute Islet Core (Edmonton, AB, Canada) and the Pathology Department of Ste-Justine Hospital (Montréal, QC, Canada) (Supplementary Table 3). Islets from non-diabetic human donors were provided by the Clinical Islet Laboratory at the University of Alberta and the National Institute of Diabetes and Digestive and Kidney Diseases – sponsored Integrated Islet Distribution Program (IIDP (RRID:SCR_014387) at City of Hope, NIH Grant # 2UC4DK098085; Supplementary Table 4). The use of human islets and pancreatic sections was approved by the Institutional Ethics Committee of the Centre Hospitalier de l’Université de Montréal (protocol no. MP-02-2019-7880).

## AUTHOR CONTRIBUTIONS

A.L.C. Conceptualization, Methodology, Investigation, Formal Analysis, Writing – Original Draft; G.M.F. Investigation; M.E. Investigation; C.T. Investigation; C.G. Investigation; M.B. Investigation; D.D.S. Investigation; J.G. Conceptualization, Validation, Writing – Review and Editing, Supervision; V.P Conceptualization, Validation, Writing – Review and Editing, Supervision, Funding Acquisition, Project Administration. V.P. is the guarantor of this work and, as such, had full access to all the data in the study and takes responsibility for the integrity of the data and the accuracy of the data analysis.

## ACKNOWLEDGEMENTS

We thank Mélanie Guévremont from the Metabolomics core facility of the CRCHUM for the β-cell mass measurements; and Dominique Gauchat and Philippe St-Onge from the Flow Cytometry core facility of the CRCHUM for assistance with measurements of β-cell proliferation. We thank Drs Pierre Bougnères, Sophie Le Fur and Marie-Pierre Belot from UMR1169, Paris Sud University for their help throughout the course of these studies.

