## Supplementary Material for "Beta-cell mass expansion during puberty involves serotonin signaling and determines glucose homeostasis in adulthood"

**CASTELL ET AL. - SUPPLEMENTARY MATERIAL**

**Supplementary Table 1**. Sources of antibodies used.

| Antibody | Dilution/Concentration | Company | Cat # |
|---|---|---|---|
| **Immunohistochemistry** | | | |
| Insulin | 1:500 | DAKO | A0564 |
| Nkx6.1 | 5μg/ml | DSHB | F55A12 |
| C-peptide | 5μg/ml | DSHB | GN-ID4 |
| Ki67 | 1:500 | Abcam | 15580 |
| BrdU | 1:1000 | BD Biosciences | 347580 |
| **Flow cytometry** | | | |
| Alexa Fluor® Mouse anti-insulin | 1:25 | BD Biosciences | 565689 |
| PE Mouse Anti-Glucagon | 1:25 | BD Biosciences | 565860 |

**Supplementary Table 2**. RT-PCR primers.

| Primer name | Forward | Reverse |
|---|---|---|
| Cyclophilin | CTTGCTGCAGACATGGTCAAC | GCCATTATGGCGTGTGAAGTC |
| GHR | GATGTTCTGAAGGGATGG | GTGGGACTGATGTTGACC |
| TPH1 | TTCTGACCTGGACTTCTGCG | GGGGTCCCCATGTTTGTAGT |
| HTR2B | AATGTCCTTGGCGGTGGCTGA | GCCAGTGGGAGGGGCCATGTA |
| HTR1D | TCACGCGGCGGCCATGATTG | CTGCCGCCAGAAGAGCGGTG |

**Supplementary Table 3**. Clinical characteristics of human pancreas donors.

| Gender | Age (y) | Weight (kgs)/BMI | Tanner stage (I-V) | Origin |
|---|---|---|---|---|
| Male | 11 | 35/16.9 | I | Ste-Justine |
| Male | 10 | 31,2/NR | I | Ste-Justine |
| Male | 9 | NR/NR | I | Ste-Justine |
| Male | 12 | NR/20.80 | NR | Alberta |
| Male | 14 | NR/21.50 | NR | Alberta |
| Female | 8.8 | 19,9/NR | I | Ste-Justine |
| Female | 14 | NR/NR | IV | Ste-Justine |
| Female | 11 | NR/NR | II | Ste-Justine |
| Female | 13 | 55.8/23.2 | IV | Ste-Justine |
| Female | 14 | NR/NR | IV | Ste-Justine |
| Female | 8 | NR/15.90 | NR | Alberta |
| Female | 10 | NR/16.80 | NR | Alberta |
| Female | 15 | NR/23.40 | NR | Alberta |

NR= not reported

**Supplementary Table 4.** Human islet donors used in study.

| | | | | | | |
|---|---|---|---|---|---|---|
| **Donors** | 1 | 2 | 3 | 4 | 5 | 6 |
| Unique identifier | SAMN13972304 | SAMN14331402 | SAMN15770453 | SAMN15877725 | SAMN15944113 | H2330 |
| Donor Age (years) | 49 | 60 | 48 | 31 | 42 | 49 |
| Donor Sex (M/F) | M | F | F | M | F | F |
| Donor BMI (kg/m$^2$) | 34.8 | 26.2 | 30.9 | 27.4 | 23.5 | 27,2 |
| Donor HbA1c | 5.5 | 5.1 | 5.8 | 5.6 | 5.4 | Not reported |
| Origin | IIDP | IIDP | IIDP | IIDP | IIDP | Clinical Islet Lab. |
| Islet isolation center | Southern California Islet Cell Resource Center | The Scharp-Lacy Research Institute | University of Wisconsin | University of Wisconsin | The Scharp-Lacy Research Institute | University of Alberta |
| Donor history of diabetes? | No | No | No | No | No | No |
| Cause of death | Cerebrovascular/ stroke | Cerebrovascular/ stroke | Cerebrovascular/ stroke | Head trauma | Cerebrovascular/ stroke | Not reported |
| Warm ischemia time (h) | Not Reported | Not Reported | Unknown | Not Reported | Not Reported | Not reported |
| Cold ischemia time (h) | 6.3 | 10.6 | 5.5 | 7.8 | 11.3 | Not reported |
| Estimated purity (%) | 90 | 90 | 90 | 95 | 95 | 72.5 |
| Estimated viability (% | 96 | 95 | 98 | 98 | 95 | 85.5 |
| Glucose-stimulated insulin secretion (SI) | SI (G2.8mM-G28mM)= 14.0 | SI (G2.8mM-G28mM)= 4.9 | SI (G2.8mM-G28mM)= 5.4 | SI (G2.8mM-G28mM)= 2.2 | SI (G2.8mM-G28mM)= 4.3 | Not reported |

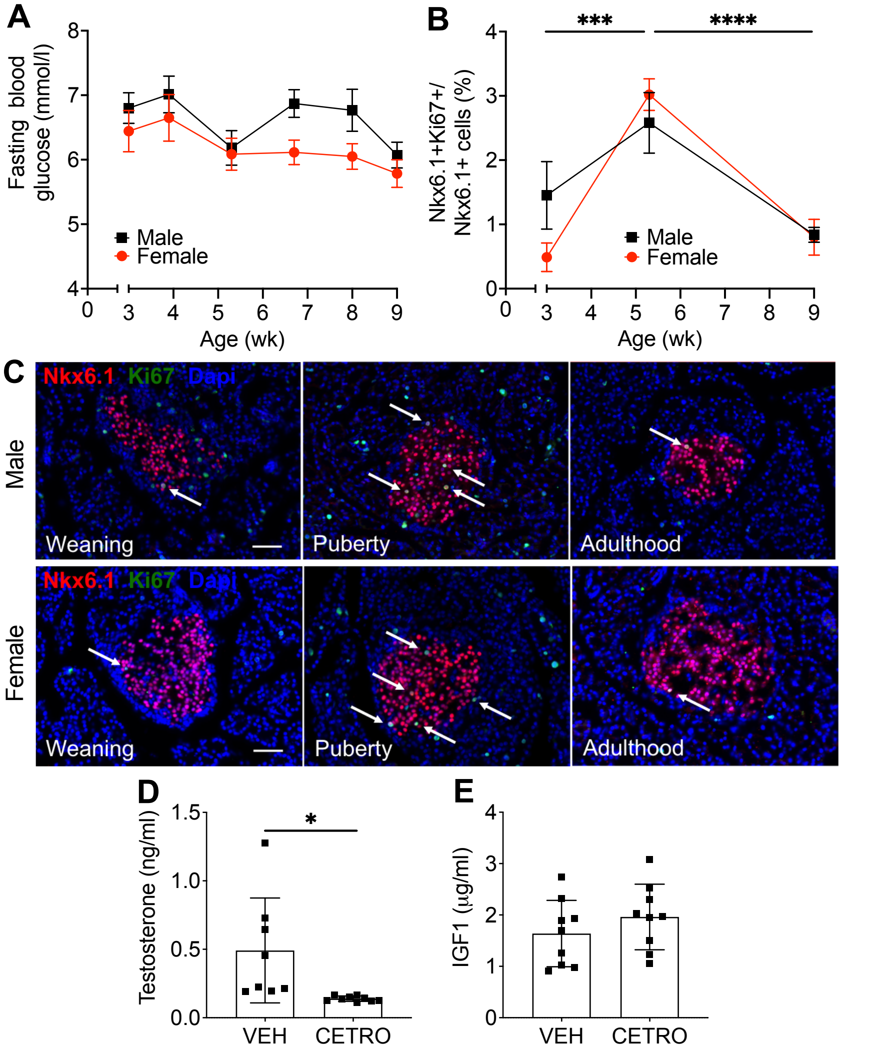


**Supplementary Fig. 1. Glucose homeostasis and β-cell proliferation in female and male rats from weaning to adulthood.** (**A**) Fasting blood glucose levels in male (black square) and female (red circle) rats from 3-9 wk of age (n=6-9). (**B-C**) β-cell proliferation as assessed by immunofluorescent staining of pancreatic sections for Ki67 and Nkx6.1 in rats at weaning (3 wk-old), puberty (~5 wk-old) or young adulthood (9 wk-old). (**B**) β-cell proliferation as a percentage of $Ki67^{+}Nkx6.1^{+}$ cells over $Nkx6.1^{+}$ cells in males (black square) and females (red circle) (n=6-8). (**C**) Representative sections showing Nkx6.1 (red), Ki67 (green) and nuclei (Dapi, blue) in male (top) and female (bottom) rats. Arrows show positive nuclei for Ki67. Scale bars, 50μm. Plasma testosterone (**D**) and IGF1 (**E**) levels at D38 in rats treated with Cetrorelix (CETRO; 100μg/d) or vehicle (VEH) from D25 to D37. Data are expressed as mean +/- SEM. ***p<0.005, ****p<0.001 following one-way ANOVA with Tukey's multiple comparisons test (**B**) or following unpaired Student's t-test (**D-E**).

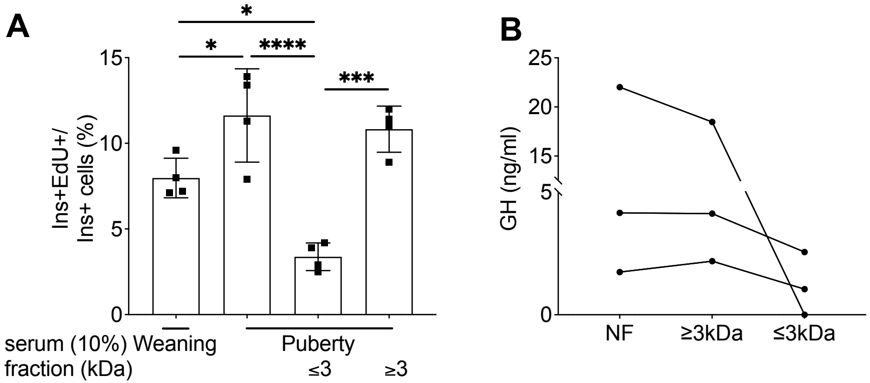


**Supplementary Fig. 2. Beta-cell proliferation in response to pubertal serum fractions.** (**A**) Male rat islets isolated at puberty were exposed to 10% weaning (3 wk-old), pubertal (~5 wk-old) or pubertal rat serum fractions ≤ or ≥ 3kDa for 72h and β-cell proliferation assessed by flow cytometry following staining for EdU and Ins and presented as a percentage of $EdU^{+}Ins^{+}$ over $Ins^{+}$ cells (n=4). (**B**) GH levels in non-fractionated (NF) pubertal rat serum or in eluate ≤ or ≥ 3 kDa as indicated (n=3). Data are expressed as mean +/- SEM. *p<0.05, ***p<0.005, ****p<0.001 following one-way ANOVA with Tukey's multiple comparisons test (**A**).

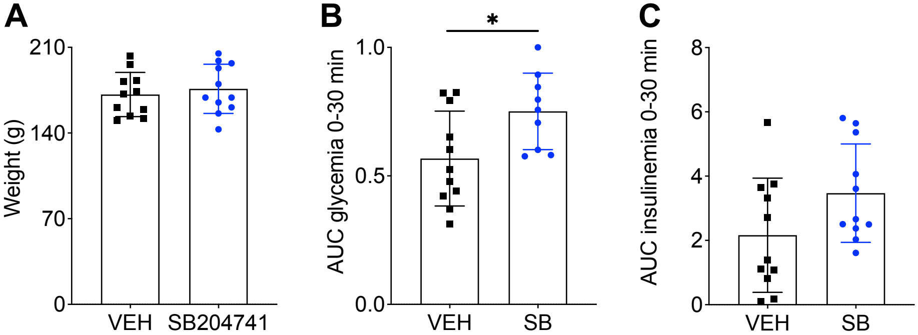


**Supplementary Fig. 3. Metabolic parameters of male rats treated with SB204741 during puberty.** Male rats were IP injected with SB204741 (SB, 1mg/kg/d) (black square) or vehicle (VEH, blue circles) from D25 to D37. Body weight (**A**), AUC of glycemia (**B**) and insulinemia (**C**) after an IP dextrose load (1g/kg) at D38. Data are expressed as mean +/- SEM. *p<0.05 following unpaired Student's t-test.

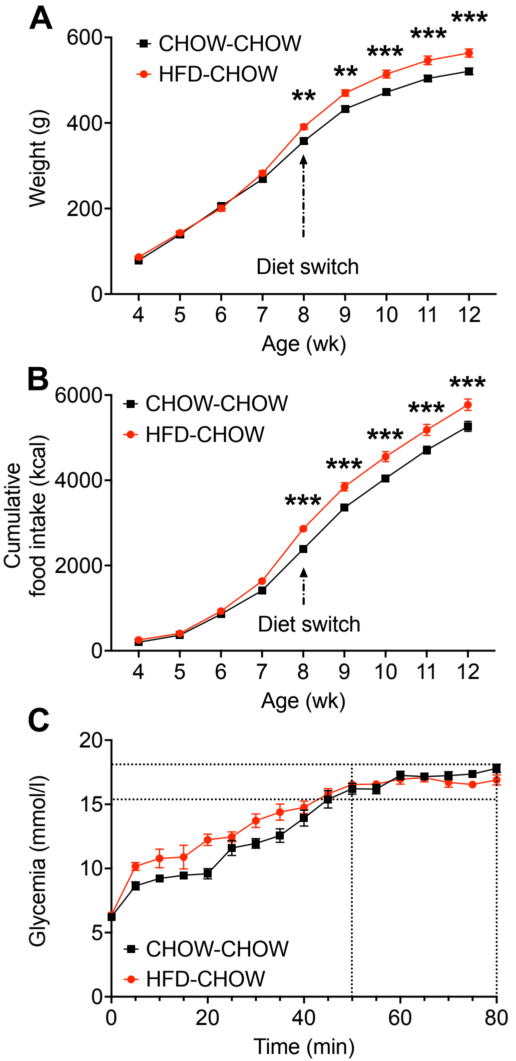


**Supplementary Fig. 4. Metabolic parameters in male rats fed a chow or HFD during puberty.** Male rats were fed a HFD (HFD, red) or a chow diet (CHOW, black) from 4 to 8 wk of age followed by a switch to a chow diet (HFD-CHOW, red; CHOW-CHOW, black) until 12 wk of age. Body weight (**A**) and cumulative food intake (**B**) during the study. (**C**) Glycemia during the HGC in HFD-CHOW and CHOW-CHOW fed animals at 12 wk of age (n=8-9). Data are expressed as mean +/- SEM. **p<0.01, ***p<0.005, following one-way ANOVA with Tukey's multiple comparisons test (**A-B**).
